# Choline degradation in *Paracoccus denitrificans*: identification of sources of formaldehyde

**DOI:** 10.1101/2023.12.12.567942

**Authors:** Trusha Parekh, Marcus Tsai, Stephen Spiro

## Abstract

*Paracoccus denitrificans* is a facultative methylotroph that can grow on methanol and methylamine as sole sources of carbon and energy. Both are oxidized to formaldehyde and then to formate, so growth on C1 substrates induces the expression of genes encoding enzymes required for the oxidation of formaldehyde and formate. This induction involves a histidine kinase response regulator pair (FlhSR) that is likely triggered by formaldehyde. Catabolism of some complex organic substrates (for example choline and L-proline betaine) also generates formaldehyde. Thus, *flhS* and *flhR* mutants that fail to induce expression of the formaldehyde catabolic enzymes cannot grow on methanol, methylamine and choline. Choline is oxidized to glycine via glycine betaine, dimethylglycine and sarcosine. By exploring *flhSR* growth phenotypes and the activities of a promoter and enzyme known to be up-regulated by formaldehyde, we identify the oxidative demethylations of glycine betaine, dimethylglycine and sarcosine as sources of formaldehyde. Growth on glycine betaine, dimethylglycine and sarcosine is accompanied by the production of up to three, two and one equivalents of formaldehyde, respectively. Genetic evidence implicates two orthologous monooxygenases in the oxidation of glycine betaine. Interestingly, one of these appears to be a bifunctional enzyme that also oxidizes L-proline betaine (stachydrine). We present preliminary evidence to suggest that growth on L-proline betaine induces expression of a formaldehyde dehydrogenase distinct from the enzyme induced during growth on other formaldehyde-generating substrates.

**IMPORTANCE:** The bacterial degradation of one carbon compounds (methanol and methylamine) and of some complex multi-carbon compounds (for example, choline) generates formaldehyde. Formaldehyde is toxic and must be removed, which can be done by oxidation to formate and then to carbon dioxide. These oxidations provide a source of energy, in some species the CO_2_ thus generated can be assimilated into biomass. Using the Gram-negative bacterium *Paracoccus denitrificans* as the experimental model, we infer that oxidation of choline to glycine generates up to three equivalents of formaldehyde and we identify the three steps in the catabolic pathway that are responsible. Our work sheds further light on metabolic pathways that are likely important in a variety of environmental contexts.

## INTRODUCTION

*Paracoccus denitrificans* is a metabolically versatile Alphaproteobacterium that is frequently detected in terrestrial and aquatic environments, in waste treatment reactors, and in association with animals and plants (1–5). The organism can grow on C1 compounds (methanol and methylamine) as sole sources of carbon and energy and has been used as a model for studies of the genetics and biochemistry of methylotrophy (6, 7). Both methanol and methylamine are oxidized to formaldehyde by PQQ-dependent periplasmic enzymes. The formaldehyde is internalized, probably by a specific transport system (8), and is oxidized to formate by a cytoplasmic glutathione (GSH) dependent formaldehyde dehydrogenase, GD-FALDH (7, 9). *P. denitrificans* also has a gene (Pden_1186, *hpbM*) encoding a predicted GSH-independent formaldehyde dehydrogenase (10, 11). Formate is oxidized to carbon dioxide, which is assimilated by the Calvin Cycle (11, 12), although other species in the *Paracoccus* genus may use the serine cycle (13).

Transcription of the genes required for the oxidation of methanol, methylamine and formaldehyde is regulated by two two-component systems comprising histidine kinases (FlhS and MoxY) and their cognate response regulators (FlhR and MoxX, respectively). MoxXY (also called MxaXY) are required for transcription of the genes encoding methanol dehydrogenase and associated proteins and *moxXY* mutants have a growth defect on methanol but not methylamine (14). FlhSR-deficient mutants are unable to grow on both methanol and methylamine (15) and the FlhSR proteins are believed to activate transcription of the operons encoding the methanol, methylamine and formaldehyde dehydrogenases, and MoxXY (7). The signal that triggers the FlhSR signal transduction pathway has not been identified biochemically, but it is suggested that both FlhS and MoxX respond to formaldehyde (7). The DeepTMHMM algorithm (16) predicts FlhS to be located in the cytoplasm, which is therefore the likely site of formaldehyde sensing.

FlhSR-deficient mutants are also unable to grow on choline as a source of carbon and energy (15). The explanation for this phenotype is that the oxidation of choline to glycine is accompanied by the production of formaldehyde and the *flhSR* mutants are poisoned because they are unable to up-regulate expression of the formaldehyde dehydrogenase. Thus, ectopic expression of the GD-FALDH rescues the choline growth defect of *flhS* and *flhR* mutants (15). Choline catabolism in *P. denitrificans* has been variously suggested to be accompanied by the production of ‘several’ (17), two (18) or three (19) equivalents of formaldehyde.

Under aerobic conditions, choline is degraded in the cytoplasm through betaine aldehyde, glycine betaine, dimethylglycine and sarcosine to glycine (Fig. 1; 20). Each of the three steps from glycine betaine to glycine involves the removal of one methyl group and is a potential source of formaldehyde. Glycine betaine can be oxidatively demethylated by a glycine betaine monooxygenase that generates dimethylglycine and formaldehyde (21, 22). In other species, a glycine betaine:homocysteine methyl transferase donates the methyl group to homocysteine and so makes methionine rather than formaldehyde (23, 24). Dimethylglycine can be oxidatively demethylated by a dimethylglycine dehydrogenase (or oxidase). These are tetrahydrofolate (THF) linked enzymes that generate methylene-THF (rather than formaldehyde) as the reaction product (25). There is also a dimethylglycine demethylase that has not been characterized biochemically but is a potential source of formaldehyde (20, 21). Sarcosine is oxidized either by a monomeric or a tetrameric sarcosine oxidase. The former generates formaldehyde as the reaction product, while the tetrameric sarcosine oxidase is THF-dependent and so makes methylene-THF (26). The THF-dependent dimethylglycine and sarcosine oxidases can make formaldehyde as a reaction product in vitro in the absence of THF (25, 26); whether these enzymes are a source of formaldehyde in vivo is not known. Depending upon the combination of enzymes used and whether or not the THF-dependent enzymes are sources of formaldehyde in vivo, the conversion of choline to glycine may be accompanied by the formation of between zero and three equivalents of formaldehyde (20, 24). *P. denitrificans* also has a characterized pathway for the degradation of proline betaine which is predicted to generate two equivalents of formaldehyde through the action of proline betaine and methyl proline demethylases (10).

**FIGURE 1.**
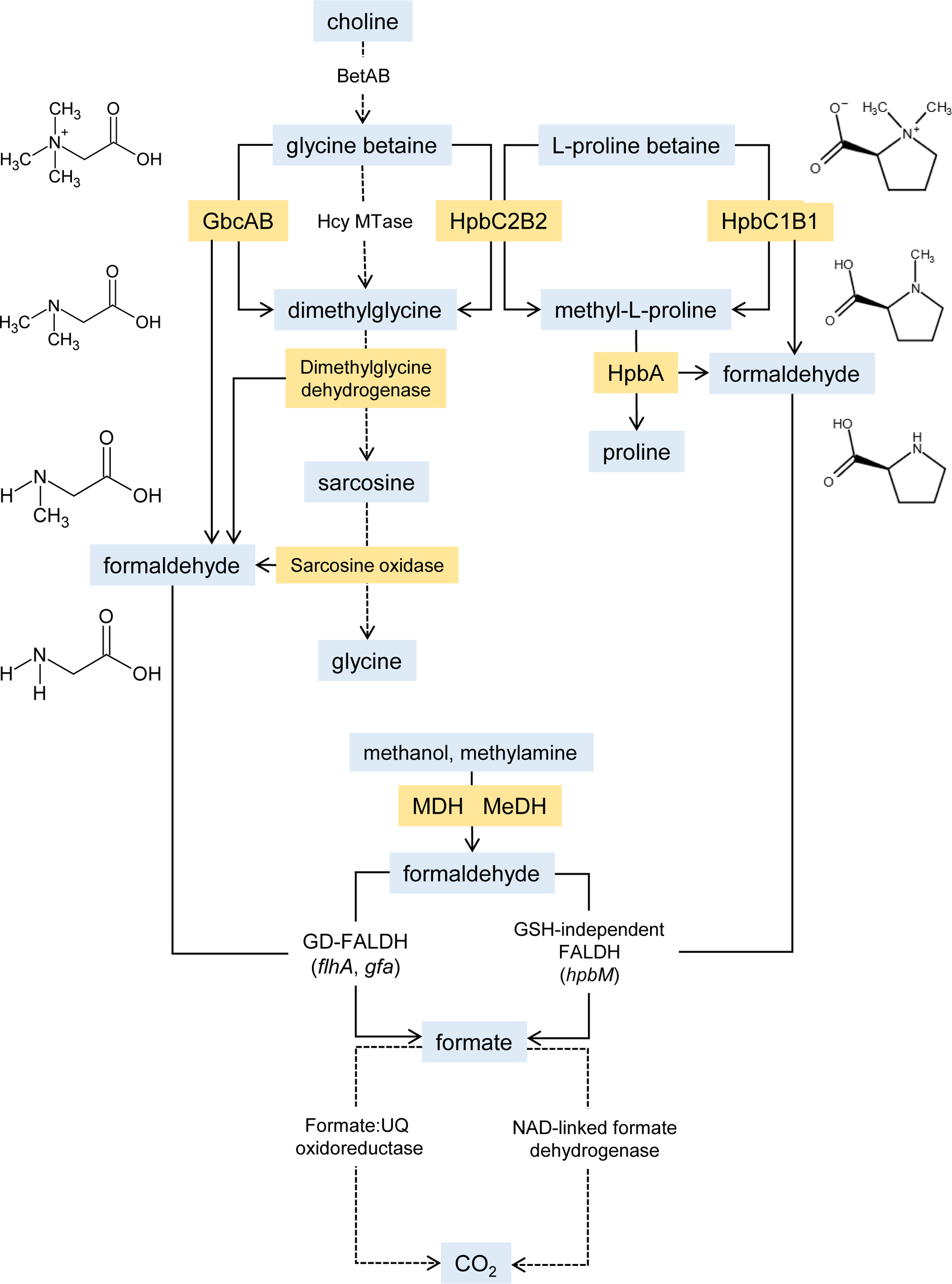
Pathways for catabolism of choline, stachydrine (L-proline betaine) and C1 substrates in *Paracoccus denitrificans*. Metabolic pathways are proposed based on previous literature (see citations in the text), the KEGG database (28) and results presented in this paper. Enzymes that are known or likely sources of formaldehyde are highlighted in orange. Solid arrows denote experimentally supported reactions, dashed lines are predictions based on the genome annotation. Based on the data presented in this paper, we suggest that formaldehyde generated from glycine betaine and dimethylglycine is oxidized predominantly by GD-FALDH, while the GSH-independent enzyme oxidizes formaldehyde generated from L-proline betaine and methyl-L-proline. The genes encoding the enzymes shown in this diagram are depicted in Figure S1.

While studying the regulation of formaldehyde metabolism in *P. denitrificans*, we became interested to identify the source(s) of the formaldehyde generated during choline catabolism. Reconstructing the pathway from the genome sequence is challenging for several reasons: (1) for some steps there are multiple candidate enzymes encoded in the genome; (2) some potentially relevant genes are duplicated, and (3) definitively assigning biochemical functions to gene products is not always possible. Therefore, we have adopted an empirical approach (informed by the genome sequence) to determine the likely pathway for choline degradation in *P. denitrificans* and the source(s) of formaldehyde. To do so, we have taken advantage of the fact that *flhSR* mutants have a defect for growth on substrates that generate formaldehyde. We present evidence to indicate that the major sources of formaldehyde are two orthologous monooxygenases that oxidatively demethylate glycine betaine to form dimethylglycine and the subsequent oxidative demethylations of both dimethylglycine and sarcosine.

## RESULTS

### Formaldehyde is a by-product of the oxidation of glycine betaine and dimethylglycine

It has been reported previously that *flhS* and *flhR* mutants are unable to grow on solid media containing choline as the sole source of carbon and energy (15). We have reconstructed unmarked deletions in *flhS* and *flhR* and confirmed that these mutants are unable to grow on solid minimal medium containing choline (Fig. S4). In liquid media, the *flhS* and *flhR* mutants grow normally to an optical density of around 0.3 before arresting (Fig. 2). This pattern is consistent with the accumulation of a toxic intermediate (likely to be formaldehyde).

**FIGURE 2.**
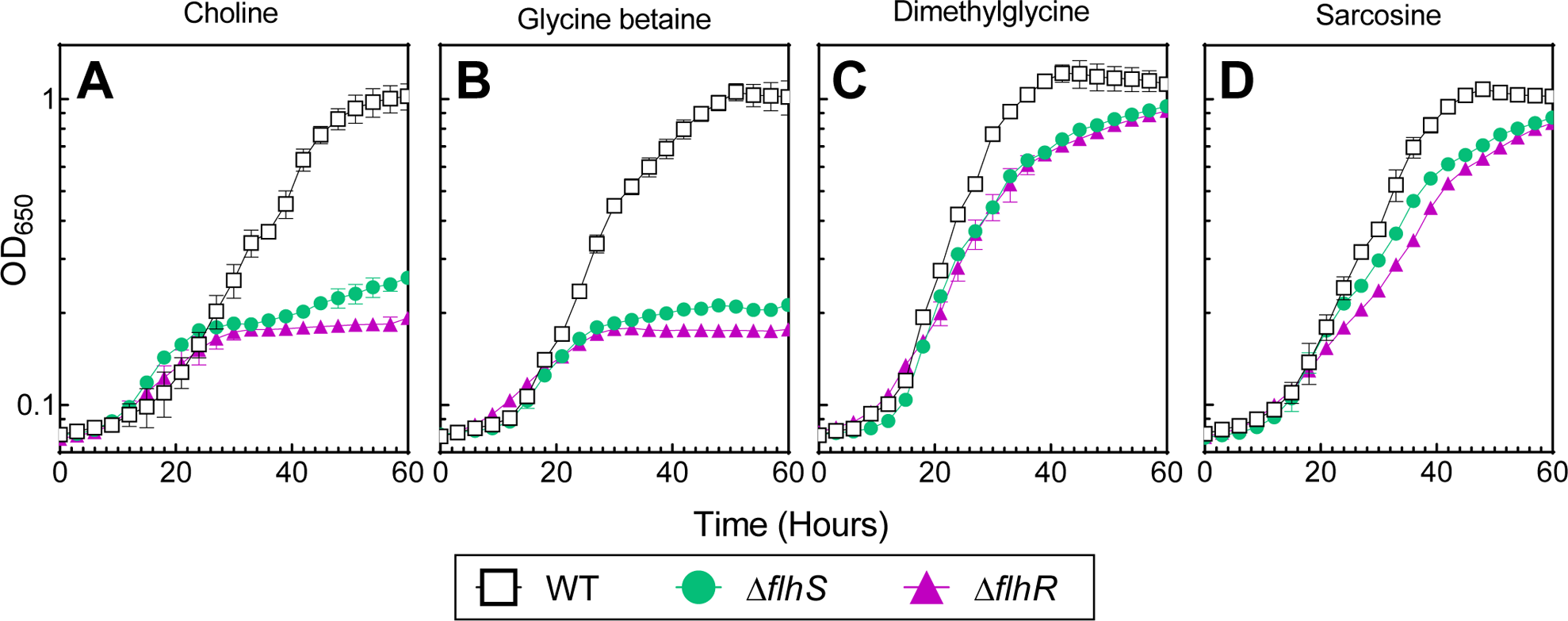
Growth phenotypes of Δ*flhS* and Δ*flhR* mutants. Wild type *P. denitrificans* and Δ*flhS* and Δ*flhR* deletion mutants were grown in minimal medium containing either (A) choline, (B) glycine betaine, (C) dimethylglycine or (D) sarcosine as sole sources of carbon and energy (substrate concentrations were adjusted such that carbon was equimolar). Growth data are the mean of three biological replicates and the error bars represent standard deviations.

We established that *P. denitrificans* grows well on intermediates of the choline degradation pathway: glycine betaine, dimethylglycine and sarcosine (Fig. 2, Table 1). In liquid media, the *flhS* and *flhR* mutants show growth defects on glycine betaine similar to the choline phenotypes (Fig. 2), whereas both mutants grow normally on dimethylglycine and sarcosine. The patterns are similar on solid media, except that the *flhS* and *flhR* mutants exhibit a partial defect for growth with dimethylglycine as the substrate (Figs. S5-7). We could detect significant concentrations of formaldehyde in the supernatants of *flhS* and *flhR* mutants grown on choline, glycine betaine and dimethylglycine (Table 1). Thus far, we conclude that formaldehyde is generated as a product of the oxidation of both glycine betaine and dimethylglycine, but that the amount of formaldehyde accumulated in dimethylglycine grown cells is insufficient to impose a phenotype on the *flhS* and *flhR* mutants.

**TABLE 1.** Growth phenotypes on solid media and formaldehyde accumulation in liquid culture supernatants.

|  | Growth phenotype<sup>a</sup> |  |  |  |  |  | Formaldehyde accumulation<sup>b</sup> |  |  |
| --- | --- | --- | --- | --- | --- | --- | --- | --- | --- |
|  | Wild type | *ΔflhS* | *ΔflhR* | *ΔgbcAB* | *ΔhpbC2B2* | *ΔgbcAB*<br>*ΔhpbC2B2* | Wild type | *ΔflhS* | *ΔflhR* |
| Succinate | +++ | +++ | +++ | +++ | +++ | +++ | - | - | - |
| Choline | +++ | - | + | +++ | + | - | - | ++ | ++ |
| Glycine betaine | +++ | - | + | +++ | - | - | - | ++ | ++ |
| Dimethylglycine | +++ | ++ | ++ | +++ | +++ | +++ | - | + | + |
| Sarcosine | +++ | +++ | ++ | +++ | +++ | +++ | - | - | - |
| Stachydrine | +++ | +++ | +++ | +++ | +++ | +++ | - | - | - |

All growth phenotypes on solid (Figs. S4-S7) and liquid (not shown) media associated with the *flhS* and *flhR* deletions could be complemented by expression of the corresponding genes *in trans* from recombinant plasmids.

### Formaldehyde generated from choline degradation activates the FlhSR signaling pathway

It has been proposed that formaldehyde generated during choline degradation activates transcription of genes required for formaldehyde oxidation (15). To confirm that choline-derived formaldehyde activates FlhSR and to determine the source(s) of formaldehyde, we took advantage of the fact that the promoter of the structural gene (*mxaF*) encoding the methanol dehydrogenase is activated by formaldehyde, by a mechanism that requires FlhSR (7, 15). We constructed an *mxaF*-*lacZ* promoter-reporter fusion and confirmed the expected low activity in cells grown on succinate and >10-fold higher activity in cells grown on choline (Table 2). The promoter fusion also had >10-fold higher activities in cells grown on glycine betaine and dimethylglycine, confirming that degradation of these substrates generates formaldehyde (Table 2). In sarcosine grown cells, *mxaF* promoter activity was intermediate between that measured in succinate and dimethylglycine grown cells, suggesting that sarcosine oxidation is also a source of formaldehyde. In all cases, induction of the *mxaF* promoter was eliminated in *flhS* and *flhR* mutants, and introduction of the corresponding genes on plasmids complemented or partially complemented the mutant phenotypes. These patterns of *mxaF* promoter activity correlate well with the formaldehyde accumulation data and mutant growth phenotypes and further suggest that degradation of glycine betaine, dimethylglycine and sarcosine produces sufficient formaldehyde to switch on an FlhSR-activated promoter.

**TABLE 2.** β-Galactosidase activities expressed from the *mxaF*-*lacZ* reporter fusion.

| | β-Galactosidase<sup>a</sup> | | | | | |
| --- | --- | --- | --- | --- | --- | --- |
| | Wild type | $\Delta flhS$ | $\Delta flhR$ | $\Delta gbcAB$ | $\Delta hpbC2B2$ | $\Delta gbcAB \Delta hpbC2B2$ |
| Succinate | 41 (2) | 39 (3) | 40 (1) | ND<sup>b</sup> | ND | ND |
| Choline | 574 (84) | 20 (17) | 30 (36) | 438 (41) | 134 (16) | 65 (9) |
| Glycine betaine | 646 (61) | 11 (9) | 2 (1) | 258 (49) | 118 (4) | 83 (5) |
| Dimethylglycine | 413 (115) | 22 (4) | 44 (33) | 319 (102) | 77 (20) | 73 (7) |
| Sarcosine | 200 (4) | 30 (25) | 51 (23) | 81 (18) | 104 (4) | 107 (2) |
| Stachydrine | 434 (101) | 16 (5) | 28 (18) | ND | ND | ND |

a. β-Galactosidase activity was measured in cells grown on the indicated carbon sources. Units of activity are as defined by Miller (38). β-Galactosidase activities are the means of triplicate assays from three independently grown cultures with the standard deviation in parentheses.
b. ND: not done.

We also measured activity of the GSH-dependent formaldehyde dehydrogenase, expression of which is believed to be activated by FlhSR (15). Similar to the pattern of *mxaF*-*lacZ* activity, GD-FALDH activity was low in extracts of cells grown on succinate, high in cells grown on choline, glycine betaine and dimethylglycine, and intermediate in cells grown on sarcosine (Table 3). Note that, if each oxidative demethylation generates one equivalent of formaldehyde, then growth on glycine betaine, dimethylglycine and sarcosine would be accompanied by production of three, two and one equivalents of formaldehyde, respectively. Accumulation of less formaldehyde is consistent with the absence of *flhSR* mutant phenotypes on dimethylglycine and sarcosine and the evidence for reduced FlhSR activity in cells grown on these substrates, compared to glycine betaine and choline. Induction of GD-FALDH activity was lost in *flhS* and *flhR* mutants (Table 3) and mutant phenotypes could be complemented by expression of the corresponding genes from a plasmid (data not shown).

**TABLE 3.** GSH-dependent formaldehyde dehydrogenase activities.

| | GSH-dependent formaldehyde dehydrogenase ($\mu$ mol NADH/min/mg protein)<sup>a</sup> | | | | | |
| --- | --- | --- | --- | --- | --- | --- |
| | Wild type | $\Delta flhS$ | $\Delta flhR$ | $\Delta gbcAB$ | $\Delta hpbC2B2$ | $\Delta gbcAB \Delta hpbC2B2$ |
| Succinate | 75 (8) | 6 (2) | 27 (13) | ND<sup>b</sup> | ND | ND |
| Choline | 361 (107) | 53 (7) | 39 (7) | 95 (35) | 264 (60) | 25 (12) |
| Glycine betaine | 539 (73) | 9 (8) | 21 (12) | 82 (36) | 207 (99) | 26 (13) |
| Dimethylglycine | 313 (62) | 37 (18) | 29 (3) | 244 (106) | 145 (49) | 53 (31) |
| Sarcosine | 135 (39) | 16 (10) | 14 (6) | 140 (90) | 32 (18) | 20 (7) |
| Stachydrine | 63 (23) | 34 (5) | 11 (7) | ND<sup>b</sup> | ND | ND |

a. GSH-dependent formaldehyde dehydrogenase activities are the means of assays from three independently grown cultures with
the standard deviation in parentheses.
b. ND: not done.

Our measurements of *mxaF* promoter activity and GD-FALDH enzyme activity both implicate the oxidation of glycine betaine, dimethylglycine and sarcosine as the sources of formaldehyde. The predicted dimethylglycine dehydrogenase encoded by Pden_1940 is likely to be required for dimethylglycine catabolism, and there are at least two candidate sarcosine oxidases encoded in the genome (Fig. S1). We focused on identifying the enzyme(s) required for growth on glycine betaine.

**TABLE 4.**
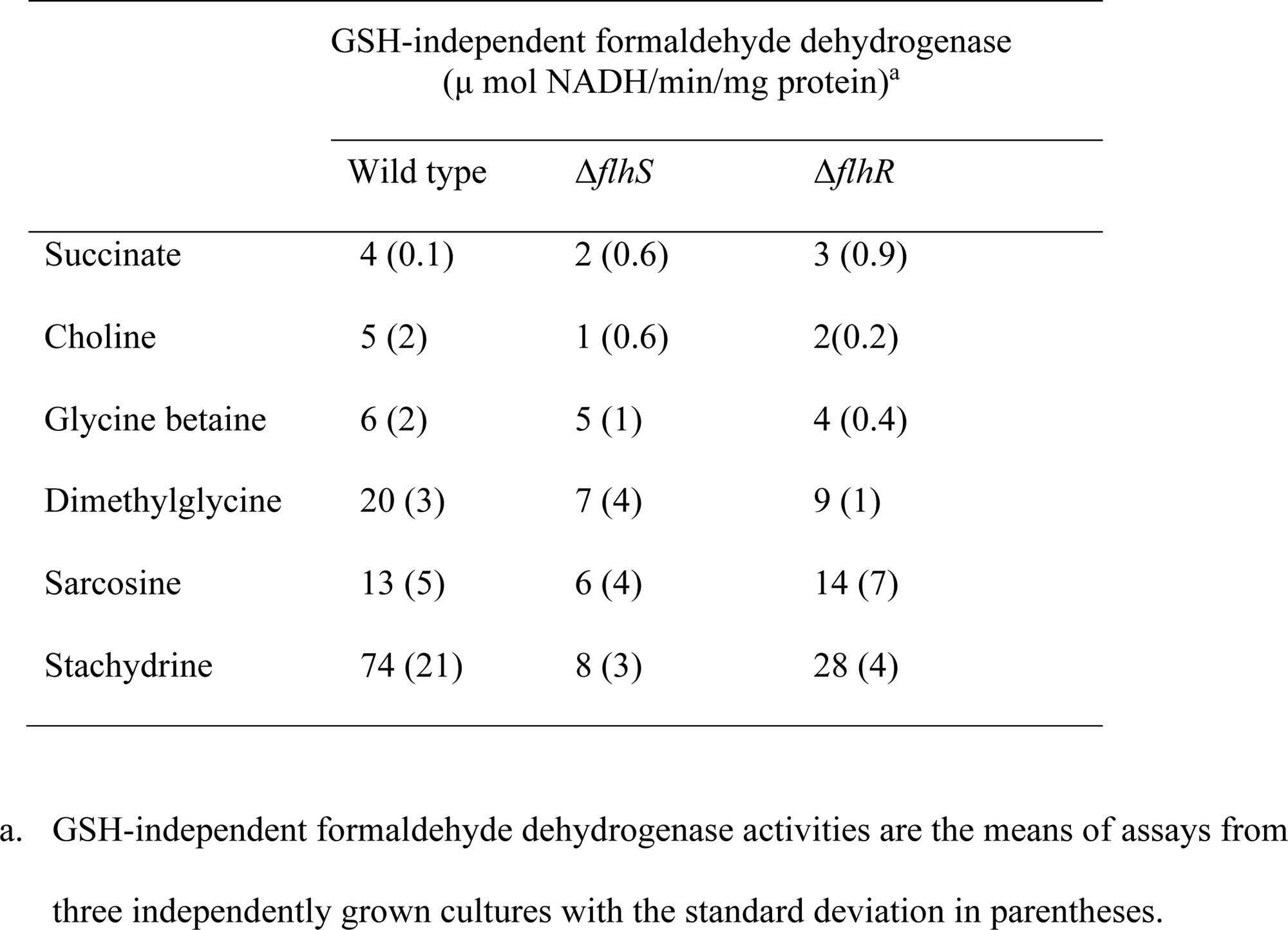
GSH-independent formaldehyde dehydrogenase activities.

| | GSH-independent formaldehyde dehydrogenase<br>($\mu$ mol NADH/min/mg protein)<sup>a</sup> | | |
| --- | --- | --- | --- |
| | Wild type | $\Delta flhS$ | $\Delta flhR$ |
| Succinate | 4 (0.1) | 2 (0.6) | 3 (0.9) |
| Choline | 5 (2) | 1 (0.6) | 2(0.2) |
| Glycine betaine | 6 (2) | 5 (1) | 4 (0.4) |
| Dimethylglycine | 20 (3) | 7 (4) | 9 (1) |
| Sarcosine | 13 (5) | 6 (4) | 14 (7) |
| Stachydrine | 74 (21) | 8 (3) | 28 (4) |

- a. GSH-independent formaldehyde dehydrogenase activities are the means of assays from three independently grown cultures with the standard deviation in parentheses.

### Overlapping functions of enzymes processing glycine betaine

In *Pseudomonas aeruginosa* and *P. syringae*, a two gene operon required for growth on glycine betaine encodes a predicted redox enzyme, GbcAB (21, 27). In *Chromohalobacter salexigens*, this enzyme has been characterized as a formaldehyde-generating glycine betaine monooxygenase (named BmoAB) comprising a Rieske non-heme iron oxygenase (BmoA) and an NADH-dependent flavin reductase, BmoB (22). In *P. denitrificans*, the predicted proteins most similar to GbcAB are encoded by Pden_4896 and 4897. We constructed a deletion mutant lacking these two genes and found only a modest defect for growth on choline and glycine betaine (Fig. 3).

**FIGURE 3.**
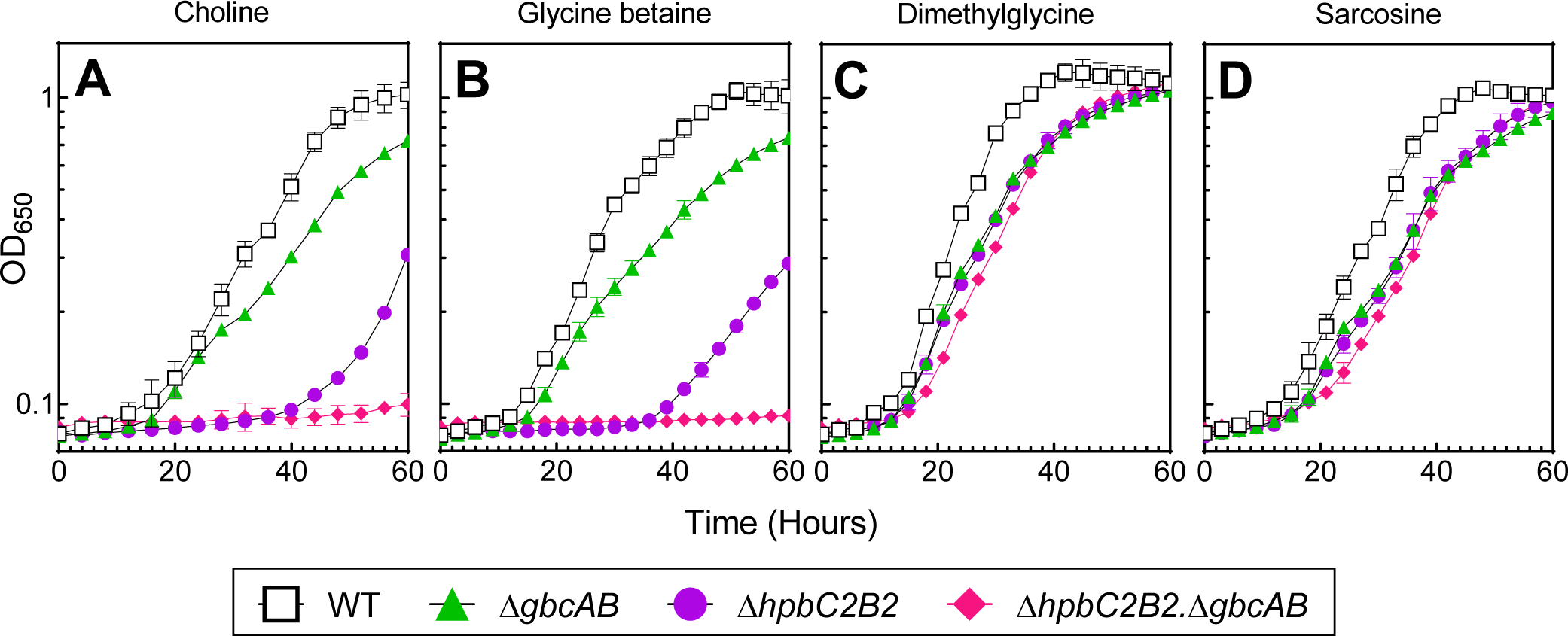
Growth phenotypes of Δ*gbcAB*, Δ*hpbC2B2* and Δ*gbcAB* Δ*hpbC2B2* mutants. Wild type *P. denitrificans* and Δ*gbcAB*, Δ*hpbC2B2* and Δ*gbcAB* Δ*hpbC2B2* mutants were grown in minimal medium containing (A) choline, (B) glycine betaine, (C) dimethylglycine, and (D) sarcosine as sole sources of carbon and energy (substrate concentrations were adjusted such that carbon was equimolar). Growth data are the mean of three biological replicates and the error bars represent standard deviations.

A previous study of stachydrine catabolism in *P. denitrificans* characterized two additional members of the GbcAB family (10). HpbC1B1 (encoded by Pden_1188-89) and HpbC2B2 (Pden_2831-32) are both annotated as stachydrine (L-proline betaine) demethylases (or monooxygenases). Deletion of both pairs of genes was required to eliminate growth on stachydrine, the implication being that both enzymes are able to oxidize stachydrine to methyl-L-proline (10). Since HpbC2B2 are more similar to GbcAB than HpbC1B1, we deleted the *hpbC2B2* (Pden_2831-32) genes and found that the mutant exhibits markedly biphasic growth when cultured on choline and glycine betaine. When subcultured into fresh medium, biphasic growth of the *hpbC2B2* mutant repeated, from which we infer that the extended lag phase and biphasic pattern is not due to the accumulation of mutations. The double mutant lacking both enzymes has a null phenotype for growth on choline and glycine betaine, implying that both GbcAB and HpbC2B2 can oxidize glycine betaine. This further suggests that HpbC2B2 has a relaxed substrate specificity being able to act on both glycine betaine (this work) and stachydrine (10).

Catabolism of stachydrine is predicted to generate two equivalents of formaldehyde (10). We found that *flhS* and *flhR* mutants have no growth defect on stachydrine (Figs. S2 and S8), as is the case for growth on dimethylglycine (Fig. 2), which we suggest also generates two equivalents of formaldehyde. From both these growth substrates, sufficient formaldehyde is made to activate the *mxaF* promoter (Table 2), but insufficient to impose a phenotype on the *flhSR* mutants (Figs. 2 and S2).

The predicted GSH-independent formaldehyde dehydrogenase is encoded by a gene (Pden_1186, *hpbM*) immediately downstream of the genes encoding the stachydrine demethylase HpbC1B1 and a proline betaine epimerase (10). We therefore speculated that expression of HpbM might be specifically induced by growth on stachydrine. We measured only background levels of GD-FALDH in extracts of cells grown on stachydrine but increased activities of a GSH-independent formaldehyde dehydrogenase compared to cells grown on all other substrates. Our results are consistent with transcriptomics data that show upregulation of *hbpM* in cells grown on stachydrine versus cells grown on methanol (10). However, the same transcriptomics data also suggest up-regulation of genes encoding the GD-FALDH by growth on stachydrine (10), which we do not see reflected in the enzyme activity.

## DISCUSSION

Considering the literature, genome annotations and metabolic predictions in the KEGG database (28), and our data, we propose a choline metabolic pathway shown in Figure 1, comprising enzymes that are the products of the genes and operons in Figure S1. Choline degradation is initiated with choline oxidation to betaine aldehyde by a choline dehydrogenase (BetA), and then to glycine betaine by a betaine aldehyde dehydrogenase (BetB). In *P. denitrificans*, these enzymes are likely encoded by Pden_1896 and Pden_1897, respectively. Upstream of these genes is Pden_1898 which is annotated to encode BetI, a repressor that likely mediates choline regulation of the *betAB* genes (29).

Our results indicate that the conversion of glycine betaine to dimethylglycine can be catalyzed primarily by HpbC2B2 (encoded by Pden_2832-2831), and in its absence, by the GbcAB (Pden_4897-4896). This step is a major source of formaldehyde in the choline degradation pathway and FlhSR signaling is required for expression of the enzyme(s) responsible for further catabolism of formaldehyde.

Conversion of dimethylglycine to sarcosine is likely catalyzed by a THF-dependent dimethylglycine dehydrogenase (encoded by Pden_1940). This enzyme can generate formaldehyde in vitro (25) and, our evidence suggests, also in vivo. In either case, our evidence indicates that this reaction does not liberate sufficient formaldehyde to inhibit growth of the *flhSR* mutants.

The *P. denitrificans* genome does not encode a predicted monomeric sarcosine oxidase but has two sets of genes annotated to encode the subunits of a tetrameric enzyme (Pden_0512, 0514-0516, and Pden_4909-4906), with the latter likely being co-expressed with a 5,10-methylene THF dehydrogenase (Pden_4905) from a shared promoter. Although the tetrameric sarcosine oxidase is THF-linked, it can make formaldehyde as a reaction product in vitro (26), our evidence suggests that this enzyme is also a source of formaldehyde in vivo.

In this work, we have successfully used mutant phenotypes and promoter and enzyme activities as probes for the endogenous generation of formaldehyde. We infer that up to three equivalents of formaldehyde may be produced during choline degradation in *P. denitrificans*. Direct quantitation of this stoichiometry would require measurement of formaldehyde production. This is challenging in part because of the pathways for the onward catabolism of formaldehyde. Ultimately, isotope-based metabolic flux analysis and metabolic modeling (11) could be used to provide a complete description of the metabolic fate of the methyl groups abstracted from quaternary amines such as choline and stachydrine.

## METHODS

### Bacterial strains, plasmids and growth conditions

Bacterial strains and plasmids used in this study are detailed in Supplementary Table S1. *Paracoccus denitrificans* Pd1222 (30) was used as the wild type strain. *Escherichia coli* S17-1 (31) was used as the host for the introduction of plasmids into *P. denitrificans* by conjugation (32). L broth (10 g tryptone.liter^−1^, 5 g yeast extract.liter^−1^, 5 g NaCl.liter^−1^) was used for the routine propagation of *E. coli* and *P. denitrificans* at 37 °C or 30 °C, respectively.

For phenotypic assays, cultures were grown in a defined minimal medium (33) containing sodium succinate (20 mM), methanol (50 mM), choline chloride (32 mM), glycine betaine (32 mM), dimethylglycine (40 mM), sarcosine (54.3 mM), stachydrine hydrochloride (12 mM), L-proline (32 mM). Aerobic cultures were grown in 200 µl minimal medium in 96-well plates shaken at 420 rpm. Growth was assayed every 2-4 hours by measurement of light scatter at 650 nm (OD650). Growth was also assessed on minimal medium agar streak plates.

### Genetic manipulations

Plasmid pSTBlue-1 (Novagen) was used as the vector for routine DNA manipulations. Triparental matings were performed by mixing log-phase *P. denitrificans*, *E. coli* S17-1 transformed with recombinant plasmid derivatives, and *E. coli* DH5α (pRK2013) in a 2:1:1 ratio and incubating on L agar for 24 h at 30 °C. *P. denitrificans* exconjugants were selected by rifampicin and indicated antibiotic(s). For conjugation and subsequent selection salt-free L broth and a low-salt (4 g sodium chloride.liter^−1^) L agar were used.

Unmarked in-frame deletions were made by allelic replacement using pK18*mobsacB* (34). Briefly, ∼500 bp regions flanking the gene(s) to be deleted were amplified separately and then ligated together by PCR using overlapping regions added to the primers. The ligated fragment was cloned into pK18*mobsacB* then mobilized into *P. denitrificans* as described above. Selection for kanamycin resistance yielded strains in which the plasmid was integrated into the chromosome by a single homologous recombination event. Recombinants with double crossovers were then isolated on the basis of sucrose resistance and kanamycin sensitivity; deletion mutations were confirmed by PCR.

For complementation tests, genes were cloned into the broad host-range expression vector pIND4 which allows induction of expression by IPTG (35). A promoter-*lacZ* reporter fusion was constructed by cloning 300 bp upstream of the start codon of the methanol dehydrogenase structural gene *mxaF* into pMP220 (36). After triparental mating into *P. denitrificans*, selection for recombinant pIND4 and pMP220 plasmids was done using 25 µg/ml kanamycin or 4 µg/ml tetracycline respectively. PCR confirmation was done with isolated plasmids.

### Formaldehyde measurement

Formaldehyde accumulation in supernatants collected from stationary phase cultures was measured using a modified NASH assay (37). 96-well plates were centrifuged at 5,000 *g* for 5 min at 4 °C after the completion of growth measurements. Triplicate 125 µl aliquots of each supernatant were mixed with 125 µl of NASH reagent (0.3% acetic acid and 0.2 % acetylacetone added freshly to a 3% ammonium acetate solution) in a clear round bottom 96 well plate and incubated at 37 °C for 50 minutes for color development.

Formaldehyde standards ranged from 0.01 mM to 1 mM. The plate was equilibrated to room temperature and the absorbance at 412 nm was recorded. Concentrations are reported after normalization to the final culture densities.

### Enzyme assays

β-galactosidase expressed from the *mxaF* promoter-*lacZ* fusion was assayed as described by Miller (38). Cultures (three independent replicates) were grown in 96 well plates, 0.05 ml samples in 0.9 ml reaction mix were made permeable with chloroform and SDS and assayed according to the standard protocol (38), the absorbance at 420 nm was measured in 96-well plates.

Glutathione-dependent formaldehyde dehydrogenase and GSH-independent formaldehyde dehydrogenase were assayed in extracts prepared from 5-50 ml cultures. Mid- to late-log phase cultures were harvested at 6,000 *g* at 4 °C for 20 min and cells were washed twice with sodium phosphate buffer (pH 7.4). The wet cell pellet was resuspended in approximately three volumes of phosphate buffer along with 2 mM DTT and 2 mM PMSF. The cell suspension was sonicated with a cycle of 20s on/20s off for 2-5 min in an ice-water bucket. The sonicated suspension was clarified by centrifugation at 16,000 g at 4 °C for 20 min.

Enzyme activity was measured at 30 °C using a Cary UV spectrophotometer. The 3 ml reaction mixture contained 50-200 µl of cell lysate in 100 mM sodium phosphate buffer (pH 7.4) and 1 mM NAD^+^. For the GSH-independent formaldehyde dehydrogenase, assays were started by the addition of 10 mM formaldehyde. For the GD-FALDH, formaldehyde addition was followed by 20 mM GSH. Specific activity was calculated as previously described using 6.22 as the millimolar extinction coefficient of NADH at 340 nm (39). Enzyme activities were assayed in triplicate in extracts of three independently grown cultures. Protein concentrations in cell lysates were measured with the Pierce BCA assay kit.

## ACKNOWLEDGMENTS

We are grateful to Sneha Narvekar for her contributions to strain constructions.

