## Supplemental Table 1 and Figures 1-8. for "Choline degradation in *Paracoccus denitrificans*: identification of sources of formaldehyde"

**Table S1: Strains and plasmids used.**

| Strain | Genotype | Source or reference |
|---|---|---|
| *E. coli* | | |
| S17-1 | *pro res*$^-$ *mod*$^+$ *recA* derivative of EC294 with integrated RP4-2 [*Tc::Mu*][*Km::Tn7*], Tp$^R$ | (1) |
| DH5α | *supE44* Δ*lacU169* (φ80 *lacZ*ΔM15) *hsdR1 recA1 endA1 gyrA96 thi-1 relA1* | (2) |
| *P. denitrificans* | | |
| Pd1222 | Wild type | (3) |
| UTD893 | Δ*flhS* | This work |
| UTD895 | Δ*flhR* | This work |
| UTD929 | Δ*gbcAB* (Pden_4896-97) | This work |
| UTD930 | Δ*hpbC2B2* (Pden_2831-32) | This work |
| UTD931 | Δ*gbcAB* Δ*hpbC2B2* | This work |
| **Plasmid** | **Details** | **Source or reference** |
| pRK2013 | Helper plasmid for conjugations | (4) |
| pIND4 | Broad-host-range expression vector | (5) |
| pMP220 | Broad-host-range *lacZ* reporter vector | (6) |
| pK18*mobsacB* | Suicide vector for making gene deletions | (7) |
| p*mxaF*-*lacZ* | *mxaF* promoter cloned in pMP220 | This work |

### Methanol and formaldehyde catabolism

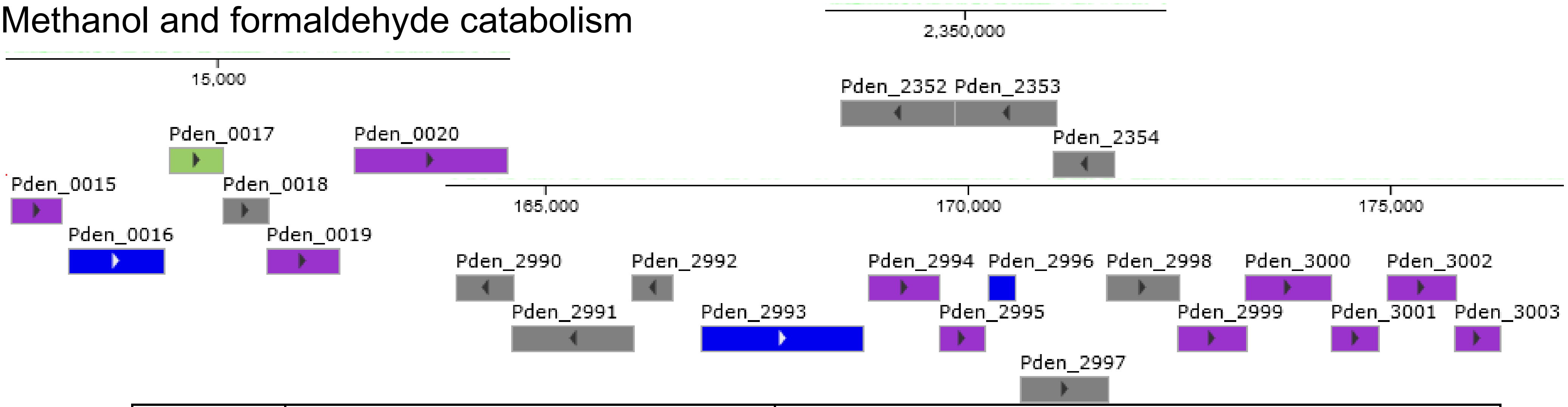


| | **Annotation (KEGG database)** | **Phenotype of mutant and reference** |
|---|---|---|
| 0015 (*gfa*) | Glutathione-dependent formaldehyde-activating protein | |
| 0016 (*flhA*) | GSH-dependent formaldehyde dehydrogenase | No growth on methanol, methylamine and choline (8) |
| 0019 (*fghA*) | Formyl glutathione hydrolase | No growth on methanol or methylamine, grows on choline (9) |
| 0020 (*xoxF*) | Lanthanide dependent methanol dehydrogenase | |
| 2352 (*flhS*) | Histidine kinase FlhS | No growth on methanol, methylamine and choline (10)<br>No growth on glycine betaine, normal growth on L-proline betaine (this paper) |
| 2354 (*flhR*) | Response regulator FlhR | No growth on methanol, methylamine and choline (10)<br>No growth on glycine betaine, normal growth on L-proline betaine (this paper) |
| 2990 (*mxaX*) | Response regulator MxaX | No growth on methanol (11) |
| 2991 (*mxaY*) | Histidine kinase MxaY | No growth on methanol (11) |
| 2993 (*mxaF*) | Methanol dehydrogenase α subunit | (12) |
| 2995 (*mxaG*) | Cytochrome $c_{551i}$ | (13) |
| 2996 (*mxaI*) | Methanol dehydrogenase β subunit | |
| 2999-3000, 3002-03 | *mxaACK…mxaLD* | |

### Choline and glycine betaine catabolism

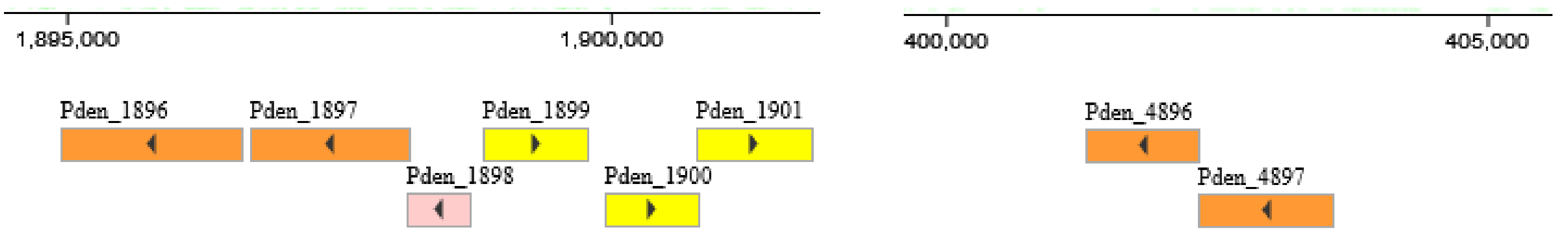


| | **Annotation (KEGG database)** | **Phenotype of mutant and reference** |
|---|---|---|
| 1896 (*betA*) | Choline dehydrogenase | |
| 1897 (*betB*) | Betaine aldehyde dehydrogenase | |
| 1898 (*betI*) | TetR family regulator | |
| 1899-1901 | ABC transporter (glycine betaine/L proline) | |
| 1940 (not shown) | $H_4F$ dependent dimethylglycine dehydrogenase | |
| 4896-97 (*gbcAB*) | Glycine betaine monooxygenase | No growth on choline or glycine betaine, when deleted with *hpbC2*/*hpbB2* (this paper) |

### Proline betaine catabolism

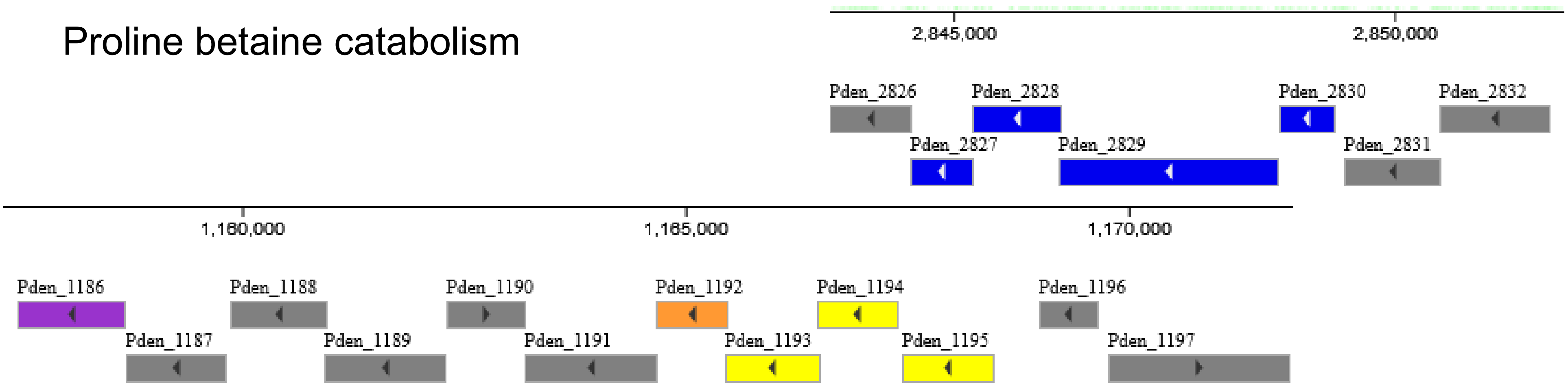


| | Annotation (KEGG database) | Phenotype of mutant and reference |
|---|---|---|
| 1186 (*hpbM*) | GSH-independent formaldehyde dehydrogenase | |
| 1187 (*hpbD*) | Proline betaine epimerase | No growth on D-proline betaine (14) |
| 1188-89 (*hpbC1*, *hpbB1*) | Stachydrine demethylase | No growth on proline betaine, when deleted with *hpbC2*/*hpbB2* (14) |
| 1190 (*hpbR*) | LysR-type regulator | |
| 1191 | FAD dependent oxidoreductase | |
| 1192 | Pyrroline-5-carboxylate reductase | |
| 1193-95 (*hpbXYZ*) | ABC transporter (glycine betaine/L proline) | |
| 1196 | TetR type regulator | |
| 1197 (*hpbA*) | Methyl proline demethylase | No growth on proline betaine or methyl proline (14) |
| 2826-2830 | Quinone-dependent formate dehydrogenase | |
| 2831-32 (*hpbC2*, *hpbB2*) | Stachydrine demethylase | No growth on proline betaine, when deleted with *hpbC1*/*hpbB1* (14)<br>No growth on choline or glycine betaine, when deleted with *gbcAB* (this paper) |

### Sarcosine catabolism

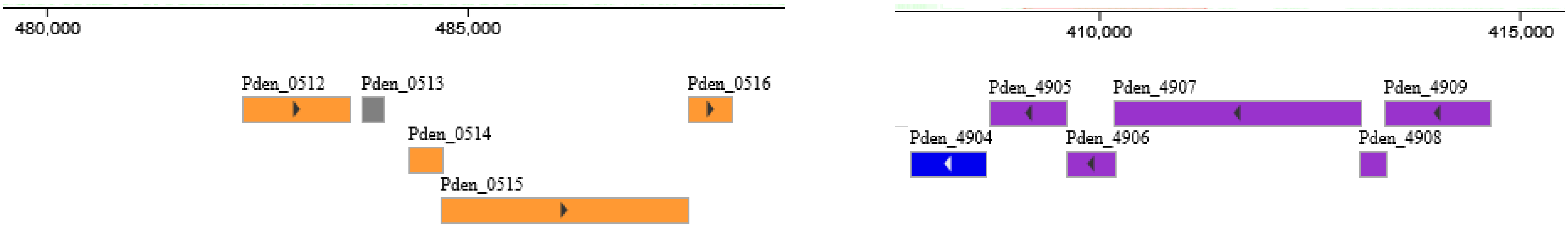


| | Annotation (KEGG database) |
|---|---|
| 0512, 0514-16 | Sarcosine oxidase |
| 4904 | Formyltetrahydrofolate deformylase |
| 4905 | Methylenetetrahydrofolate dehydrogenase |
| 4906-09 | Sarcosine oxidase |

**Figure S1:** *Paracoccus denitrificans* genes encoding enzymes and accessory proteins known or predicted to be involved in the catabolism of methanol, formaldehyde, choline, glycine betaine, proline betaine (stachydrine), and sarcosine.

Gene maps and annotations are from the KEGG database (15).

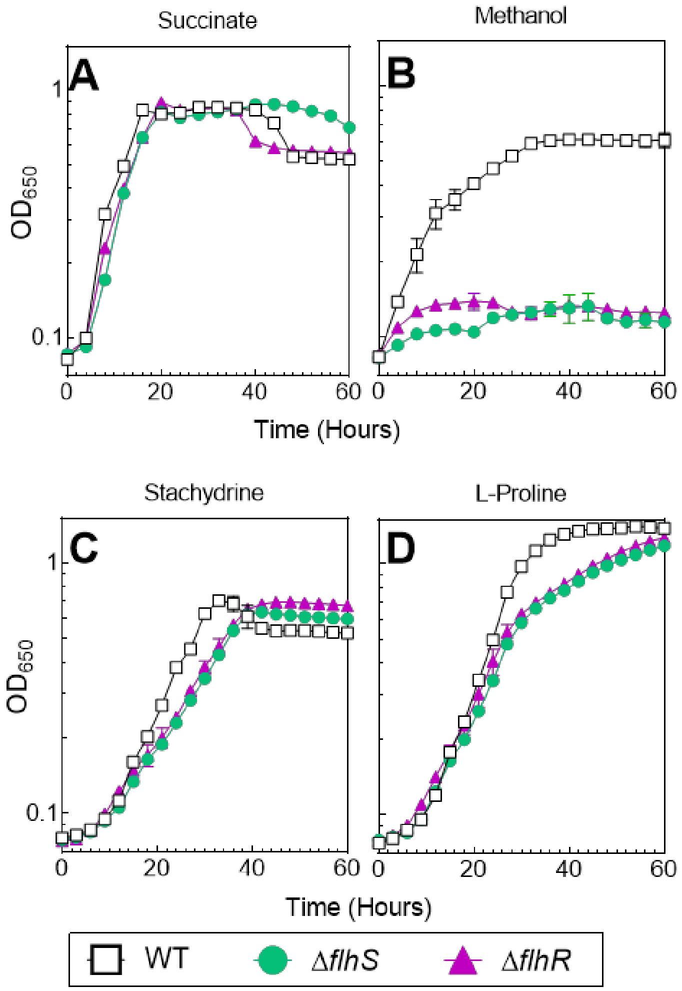


**Figure S2. Growth of Δ*flhS* and Δ*flhR* mutants on non-choline pathway substrates.**

Wild type *P. denitrificans* and Δ*flhS* and Δ*flhR* deletion mutants were grown in minimal medium containing either (A) succinate, (B) methanol, (C) stachydrine, or (D) L-proline as sole sources of carbon and energy (substrate concentrations were adjusted such that carbon was equimolar). Growth data are the mean of three biological replicates and the error bars represent standard deviations.

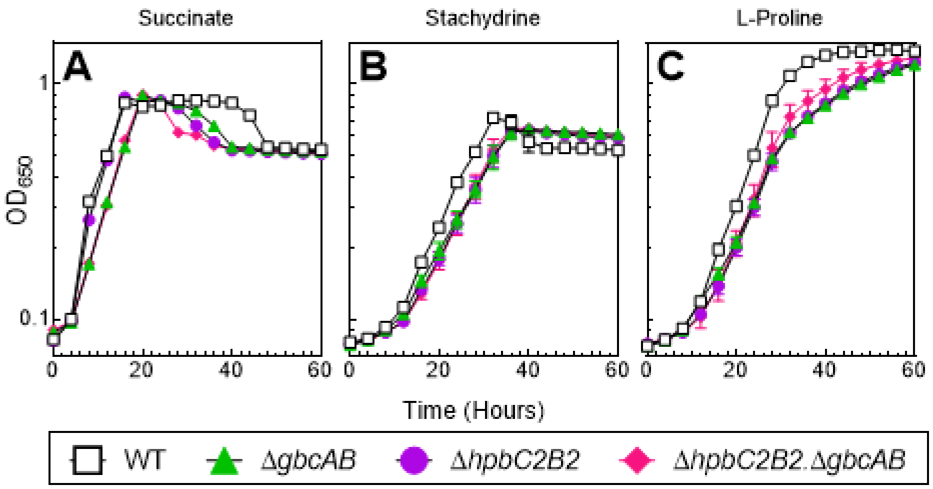


**Figure S3. Growth phenotypes of Δ*gbcAB*, Δ*hpbC2B2* and Δ*hpbC2B2* Δ*gbcAB* mutants on non-choline pathway substrates.**

Wild type *P. denitrificans* and Δ*gbcAB*, Δ*hpbC2B2* and Δ*hpbC2B2* Δ*gbcAB* mutants were grown in minimal medium containing (A) succinate, (B) stachydrine, or (C) proline as sole sources of carbon and energy (substrate concentrations were adjusted such that carbon was equimolar). Growth data are the mean of three biological replicates and the error bars represent standard deviations.

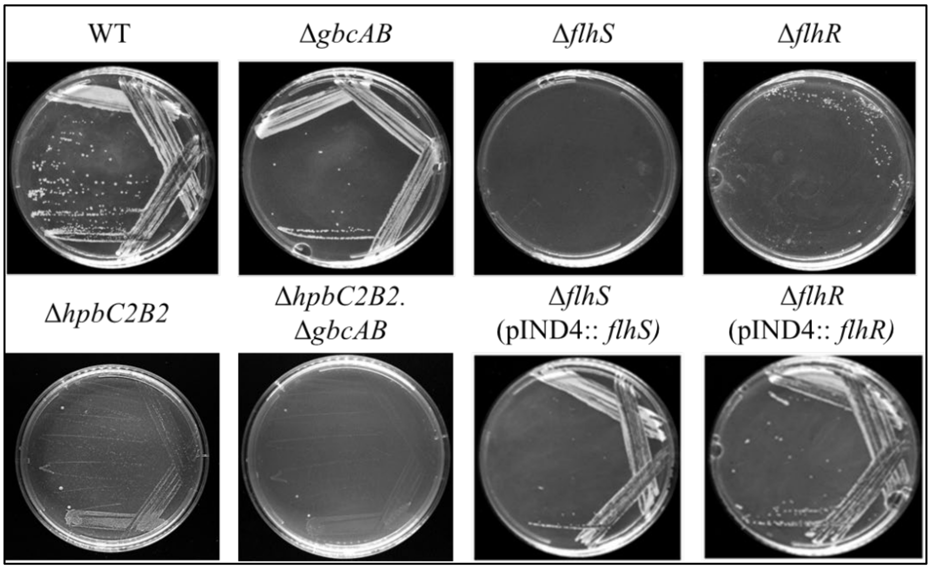


**Figure S4. Growth on solid choline minimal medium.**

The indicated strains were grown on minimal medium agar plates containing 32 mM choline. Each of the WT and mutant strains were streaked for single colonies from overnight cultures and were incubated at 30°C for 3 days. Growth in each streak was recorded from images captured with the Invitrogen iBright CL1500 Imaging system and is summarized in Table 2.

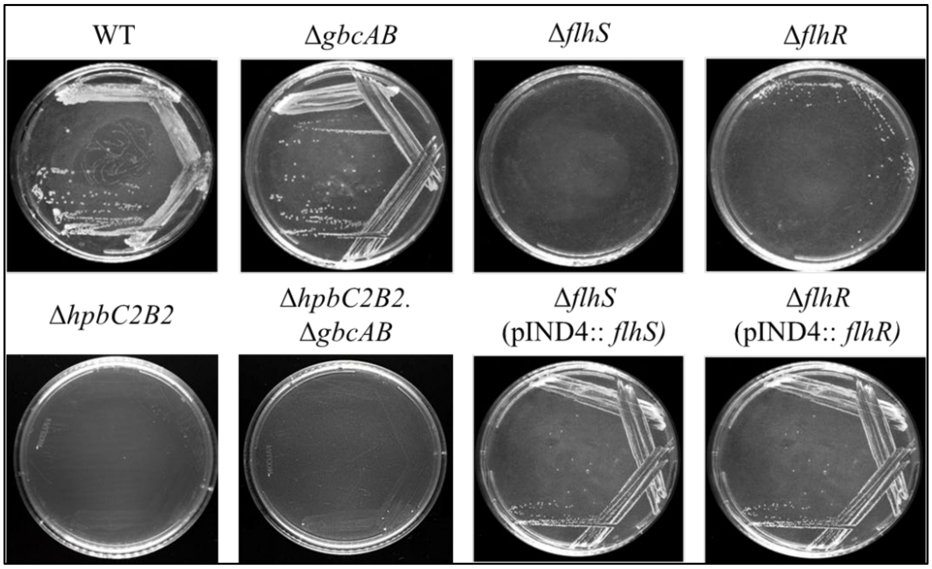


**Figure S5. Growth on solid glycine betaine minimal medium.**

The indicated strains were grown on minimal medium agar plates containing 32 mM glycine betaine. Each of the WT and mutant strains were streaked for single colonies from overnight cultures and were incubated at 30°C for 3 days. Growth in each streak was recorded from images captured with the Invitrogen iBright CL1500 Imaging system and is summarized in Table 2.

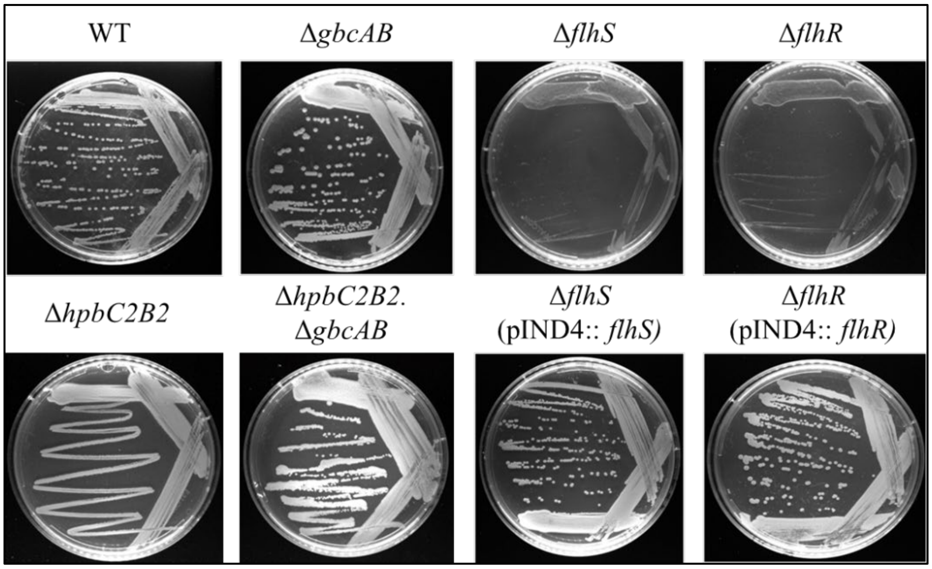


**Figure S6. Growth on solid dimethylglycine minimal medium.**

The indicated strains were grown on minimal medium agar plates containing 40 mM dimethylglycine. Each of the WT and mutant strains were streaked for single colonies from overnight cultures and were incubated at 30°C for 3 days. Growth in each streak was recorded from images captured with the Invitrogen iBright CL1500 Imaging system and is summarized in Table 2.

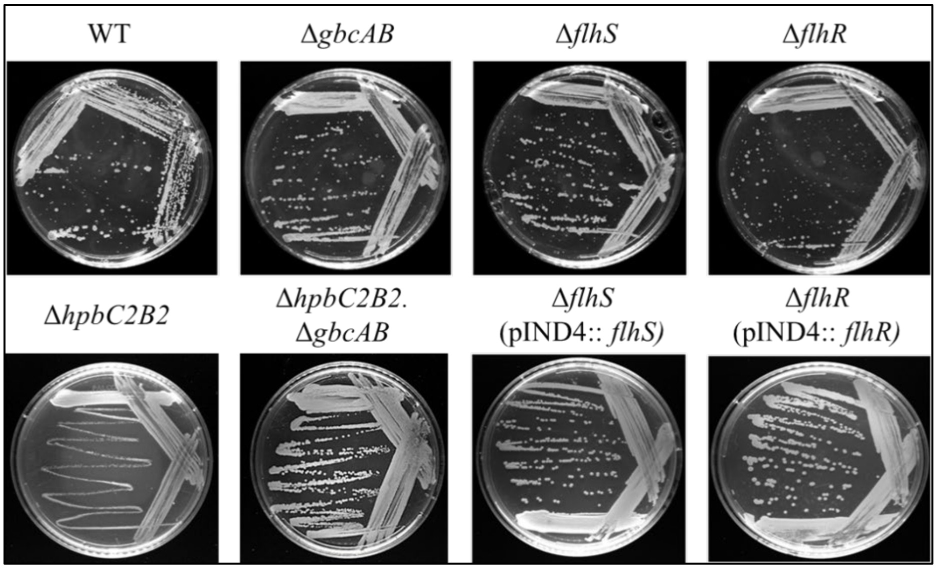


**Figure S7. Growth on solid sarcosine minimal medium.**

The indicated strains were grown on minimal medium agar plates containing 54.3 mM sarcosine. Each of the WT and mutant strains were streaked for single colonies from overnight cultures and were incubated at 30°C for 3 days. Growth in each streak was recorded from images captured with the Invitrogen iBright CL1500 Imaging system and is summarized in Table 2.

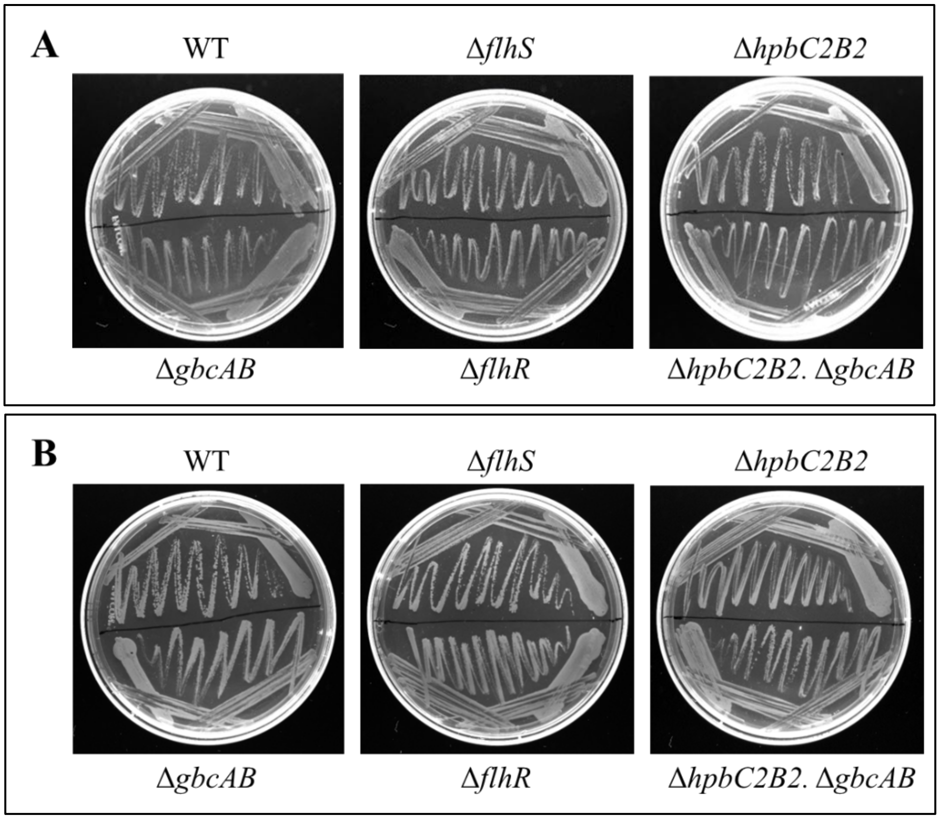


**Figure S8. Growth of WT and mutant strains on stachydrine and L-proline.**

The indicated strains were grown on solid minimal medium containing (A) 12 mM stachydrine and (B) 32 mM L-proline. Each of the WT and mutant strains were streaked for single colonies from overnight cultures and were incubated at 30°C for 3 days. Growth in each streak was recorded from images captured with the Invitrogen iBright CL1500 Imaging system and is summarized in Table 2.
